# Hierarchical autoencoder-based integration improves performance in multi-omics cancer survival models through soft modality selection

**DOI:** 10.1101/2021.09.16.460589

**Authors:** David Wissel, Daniel Rowson, Valentina Boeva

## Abstract

With decreasing costs of high-throughput sequencing, more and more datasets providing omics profiles of cancer patients become available. Thus, novel survival analysis approaches integrating these differently sized and heterogeneous molecular and clinical groups of variables start being developed. Due to the difficulty of the task of multi-omics data integration, the Cox Proportional-Hazards (PH) model using clinical data has remained one of the best-performing techniques, barely outperformed by models using molecular data modalities. There is therefore a need for methods that can successfully perform multi-omics integration in survival analysis and outperform the clinical Cox PH model. Moreover, while certain deep learning methods have been shown to provide state-of-the-art accuracy of cancer survival prediction, most of them show no benefit or even decay in performance when integrating a larger number of modalities, further motivating a need to investigate how modality-specific representations should be integrated when using neural networks for multi-omics integration. We benchmarked multiple integration techniques for a neural network architecture, revealing that hierarchical autoencoder-based integration of modality-specific representations outperformed other methods such as max-pooling and was comparable with state-of-the-art statistical approaches for multi-omics integration. Further, we showed that the hierarchical autoencoder-based integration of modality-specific representations achieved increased performance through a soft modality selection mechanism, focusing on the most informative modalities for each cancer. We thus framed multiomics integration as a partial group-wise feature selection problem, highlighting that only those models performed well that could adequately weight important modalities in the presence of the high noise imposed by less important modalities.

## 1 Introduction

Accurate prediction of survival times is essential for clinicians and researchers to decide treatment and identify which factors drive survival. The Cox Proportional-Hazards (Cox PH) model [Cox, 1972, Breslow, 1975] is still the *de facto* standard for survival analysis today, despite proposals for various other methods such as random survival forests [Ishwaran et al., 2008], boosting [Hothorn et al., 2010] and neural networks [Katzman et al., 2018].

Survival analysis of cancer patients can be particularly challenging due to the heterogeneous nature of the disease, even for patients suffering from the same type of cancer [Polyak et al., 2011, Fisher et al., 2013, Melo et al., 2013]. With the advent of high throughput sequencing technologies, researchers hoped to leverage the information inherent in molecular data such as gene expression, DNA methylation, genomic mutations, and others (jointly referred to as multi-omics) to help explain and mitigate this heterogeneity.

However, even using this wealth of newly available biological data in large scale projects such as The Cancer Genome Atlas Program (TCGA) [Tomczak et al., 2015], significant improvements in performance in cancer survival analysis as measured by discriminative performance metrics such as Harrell’s concordance [Harrell et al., 1982] (c-index) have been elusive. In particular, Herrmann et al. [2021] showed that the Cox PH model using only clinical data outperformed most other methods, even when these were designed to integrate multi-omics data. We now briefly survey related work on multi-omics survival models, before moving on to our contributions.

There have been various proposals for both statistical and neural models that perform multi-omics integration in the context of cancer survival analysis. Most statistical models for multi-omics integration rely on what we refer to as *integration through regularization*. That is, making sure that each modality gets its “fair share” by adjusting, *e*.*g*., the regularization hyperparameter specifically for each modality (in regularized Cox models) [Boulesteix et al., 2017] or up weighting the split point criterion for specific modalities (in random forests) [Hornung and Wright, 2019a] (Figure 1A). Hornung and Wright [2019a] proposed five variations of the random forest algorithm [Breiman, 2001], all of which change the split point selection by taking into account that the input variables belong to different groups (*e*.*g*., multi-omics data). BlockForest, their best performing method, statistically significantly outperformed random survival forest in their work [Hornung and Wright, 2019a] and was shown capable of outperforming the clinical Cox PH model on TCGA by Herrmann et al. [2021]. Boulesteix et al. [2017] proposed a modified Lasso regularized Cox PH model that scales the Lasso penalty *λ* with a group-specific penalty factor that can be chosen through *a priori* knowledge or using cross-validation. The authors showed that their new model, termed *IPF-Lasso*, performed better than a Lasso regularized Cox PH model in simulations and on TCGA. Klau et al. [2018] introduced a sequential Lasso regularized Cox PH model approach based on offsetting, called *prioritylasso*, which considers input modality groups one at a time and uses the previous model prediction as an unregularized offset for the model fit with the next input group. Their model outperformed a Lasso regularized Cox PH model and offers clinicians the advantage of being able to decide which groups of variables should be preferentially included in the model [Klau et al., 2018].

**Figure 1:**
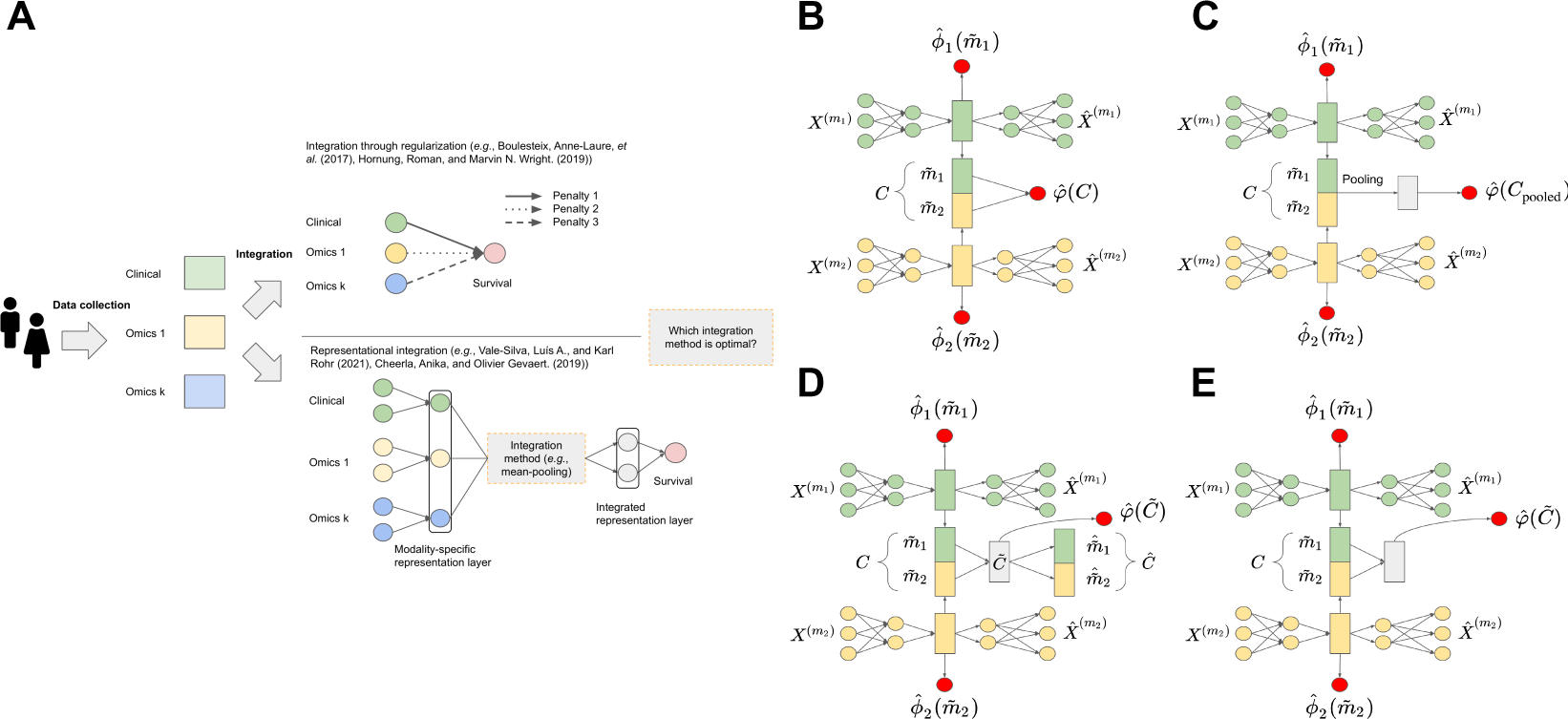
Overview of our work on establishing the best integration method when attempting multi-omics integration of representations learned by modality-specific neural networks. Two input modalities shown for simplicity in all architecture diagrams. Let 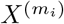 denote the submatrix of *X* containing only columns belonging to the i-th input variable group, and let 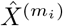 denote a reconstruction of 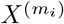. Let 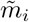 be the modality-specific representation learned based on input modality *i*. Let 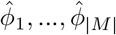 be predicted relative risks based on the modality-specific representation learned for each input modality, while 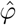 represents the predicted relative risk based on the overall joint representation. A. Pipeline of multi-omics integration in a survival context. B. Architecture diagram of ConcatSAE. C. Architecture diagram of MeanSAE and MaxSAE. Pooling may refer to either max-pooling or mean-pooling. D. Architecture diagram of HierarchicalSAE. E. Architecture diagram of HierarchicalSAE (no decoder).

Neural models for multi-omics integration generally rely on a separate network per input modality, the learned representations of which are then integrated using operations such as mean-pooling, max-pooling, or concatenation. We refer to first learning modality-specific representations which are then integrated as *representational integration* (Figure 1A). Cheerla and Gevaert [2019] proposed a new architecture to predict cancer survival by integrating gene expression, miRNA, clinical data, and whole slide images by using a similarity loss combined with mean-pooling. Their model benefitted (that is, exhibited an increased c-index) from pan-cancer training relative to training solely on each cancer type for most considered cancers. Tong et al. [2020] explored multi-modal autoencoders for the integration of multi-omics data in breast cancer survival. They proposed two architectures, each of which first fitted a dedicated autoencoder per modality. Their second architecture based on concatenating the modality-specific representations, *ConcatAE*, performed best on TCGA breast cancer when integrating DNA methylation and miRNA while using PCA for dimensionality reduction. Vale-Silva and Rohr [2021] presented a multi-modal neural network, *MultiSurv*, which used maxpooling to integrate the representations constructed by a separate neural network per modality, in addition to using a non-proportional hazards loss by assuming discrete follow-up times. *MultiSurv* outperformed a regularized Cox PH model, Random Survival Forest, *DeepSurv* [Katzman et al., 2018] and *DeepHit* [Lee et al., 2018] on single modalities and was able to achieve state-of-the-art results when integrating multiple modalities. While Vale-Silva and Rohr [2021] reported that they tried different integration techniques (such as sum-pooling, product-pooling, and concatenation among others), they state that they were not able to find consistent improvements over max-pooling, and thus did not provide results for each integration method. Zhang et al. [2021] developed a multi-task pan-cancer multi-omics model that integrated the modality-specific representations learned by one neural network per modality using one top-level autoencoder. Their model deemed *OmiEmbed*, outperformed *DeepSurv* and a Cox PH model based on the dimensionality-reduced input data in terms of the c-index and the integrated Brier score on the full TCGA dataset by integrating gene expression, DNA methylation, and miRNA.

Further work on variational autoencoder (VAE) architectures in the context of multi-omics integration by Simidjievski et al. [2019] showed that not all architectures and integration methods perform equally well. In particular, the authors considered breast cancer patients from METABRIC [Curtis et al., 2012] and compared the performance of different VAE architectures and integration techniques (among others notably a hierarchical autoencoder) on the embedding quality and three downstream classification tasks for which the learned embeddings were used (IHC subtypes, gene expression subtypes, and iCluster subtypes). The authors found that their hierarchical VAE was among the best-considered architectures. Tan et al. [2020] presented an application of supervised autoencoders on TCGA data. The authors tried to predict binarized clinical endpoints (*e*.*g*., overall survival, diseasefree survival) from TCGA data using methylation, miRNA, mRNA, and reversed-phase protein arrays (RPPA). They trained one supervised autoencoder for each modality group and integrated their modality-specific representations using mean-pooling, from which they predicted their chosen endpoints. Using this architecture, the authors outperformed other machine learning models such as random forests and support vector machines.

We analyzed published approaches and found a gap, in that, to the best of our knowledge, no previous work had analyzed the effect of different methods for representational integration as increasing numbers of modalities were considered. Instead, most previous works which analyzed the effect of different integration methods usually restricted themselves to two to three modalities or did not compare integration methods while varying the number of considered modalities. Previous work, however, has shown that many multi-omics neural models perform better when integrating only a subset of available modalities. We were thus interested in exploring an integration method that was robust to the noise imposed by adding further modalities and at least did not get significantly worse when adding further modalities.

Our study thus explored the effect of four different integration techniques, namely mean-pooling, max-pooling, concatenation, and a hierarchical autoencoder on the performance of one underlying neural architecture. Overall, our main contributions were three-fold:

1. We showed that hierarchical autoencoder-based multi-omics integration of modality-specific representations could outperform other integration techniques commonly used for this purpose, including mean-pooling, max-pooling, and concatenation.
2. We showed that neural models using hierarchical autoencoder-based integration performed on-par with or better than the best performing statistical models when integrating all available multi-omics modalities and clinical data from TCGA. To the best of our knowledge, there has been little to no work drawing this comparison, as many studies tend to focus on either neural models or statistical models.
3. We posited that, in a survival or risk prediction context, multi-omics integration can partially be framed as a group (*i*.*e*., modality) feature selection problem. In particular, the best performing multi-omics methods seem to have accurately weighted each modality and performed a soft modality selection (hence not completely excluding additional blocks, but focusing on the most informative ones, such as clinical and gene expression).

## 2 Methods

### 2.1 Architecture

We considered representational multi-omics integration, in which we, similar to other works, fitted modality-specific neural networks, which were then combined using different integration methods. For each modality-specific network, we considered identical supervised autoencoders, using a Cox loss for supervision to ensure that survival-specific representations were learned. In particular, we chose autoencoders such that we would not have to run feature selection before training.

More formally, within our models, each input modality *m*_*i*_ ∈ *M* had a separate autoencoder that compressed the features of each group 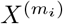 into a modality-specific representation 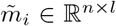, where *n* represents the number of samples. Since we modeled survival using supervised autoencoders, we also introduced layers from each modality-specific representation to a corresponding predicted relative risk 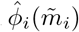. To produce a combined representation, an integration method Φ : ℝ^*l×m*^ →ℝ^*p*^ then took the latent space representations from all modality-specific autoencoders and compressed them into an integrated representation which was used for the final prediction of the relative risk for each patient 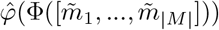 (Figure 1B-E).

We used the mean squared error (MSE) as the reconstruction loss for all autoencoders and the Breslow approximation of the negative partial log-likelihood as the loss for all predicted relative risk values [Breslow, 1975]. We regularized all layers within our autoencoders with *l*_2_ regularization, as a standard regularization choice to combat overfitting. Let *δ*_*i*_ be the event indicator for patient *i* where *δ*_*i*_ = 1 if patient *i* experienced the event during the study and *δ*_*i*_ = 0 otherwise. Further, supposing that *Y*_*i*_ is the time of the event for patient *i*, we let *T*_*i*_ = min(*Y*_*i*_, *S*_*i*_) where *S*_*i*_ is the time at which patient *i* was censored. *γ* denotes a hyperparameter controlling the strength of the reconstruction loss of each modality-specific autoencoder and *λ* denotes a hyperparameter controlling the strength of the *l*_2_ regularization. Let 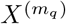 denote the submatrix of *X* containing only columns belonging to the q-th input modality and 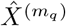 denote a reconstruction 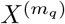. Then, the full loss for our autoencoders becomes:

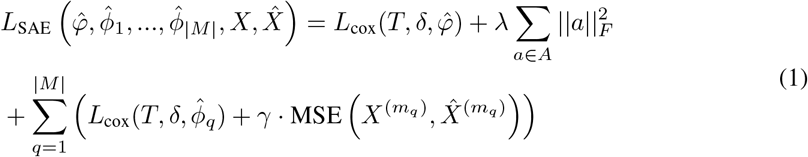

Here, ∥. ∥ _*F*_ denotes the Frobenius norm, and *L*_cox_ the Breslow approximation of the negative partial log-likelihood. *A* denotes the set of all weight matrices corresponding to each layer in our networks.

### 2.2 Integration methods

Overall, we considered the following integration methods that all took the representations learned by each modality-specific neural network and attempted to integrate them to produce or learn a joint representation amenable to survival prediction:

1. Concatenation (ConcatSAE) (Figure 1B)

2. Mean-pooling (MeanSAE) (Figure 1C)

3. Max-pooling (MaxSAE) (Figure 1C)

4. A second level autoencoder (HierarchicalSAE) (Figure 1D)

5. A second level autoencoder without the decoder (HierarchicalSAE (no decoder)) (Figure 1E), which was used primarily for ablation studies.

For HierarchicalSAE, we modified our original loss function (Eq. 1) to include the reconstruction loss of the second level autoencoder-based integration:

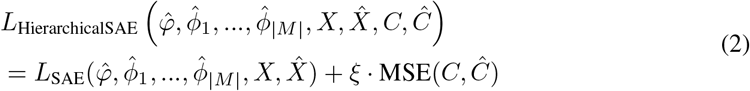

Where *ξ* represents a hyperparameter controlling the strength of the reconstruction loss of the second level autoencoder, which, identical to *γ*, was set equal to the batch size of our models.

Notably, initially, we did not include any hidden layers after each integration method, since we were interested in establishing the ability of the models to learn a survival-relevant representation without additional transformation.

### 2.3 Implementation

We developed all neural networks using Pytorch [Paszke et al., 2019] and skorch [Tietz et al., 2017]. We used batch normalization throughout the networks [Ioffe and Szegedy, 2015] and Parametric Rectified Linear Units for non-linear activations [He et al., 2015]. For all neural networks, we performed standardization of all features before fitting the model.

Training was performed using a validation set of 10% of the training set which was used for both early stopping and learning rate scheduling. We fitted all models using the Adam optimizer with an initial learning rate of 0.01 which was reduced by a factor of 10 if the validation loss did not decrease for three epochs. We trained all models for a maximum of 100 epochs using no batching, since all datasets were quite small (Table 1). Architecturally, all representations (*i*.*e*., latent spaces of the modality-specific autoencoders, as well as the latent space of HierarchicalSAE) were set to 64, and hidden layers (if architectures contained them) were fixed to 128 nodes. The *l*_2_ penalty hyperparameter *λ* was set equal to 0.001 for all models in initial experiments, the largest initial choice recommended by Smith [2018] since our datasets were quite small. We scaled the reconstruction loss by multiplying it by the batch size (*i*.*e*., *γ* = batch size).

**Table 1:**
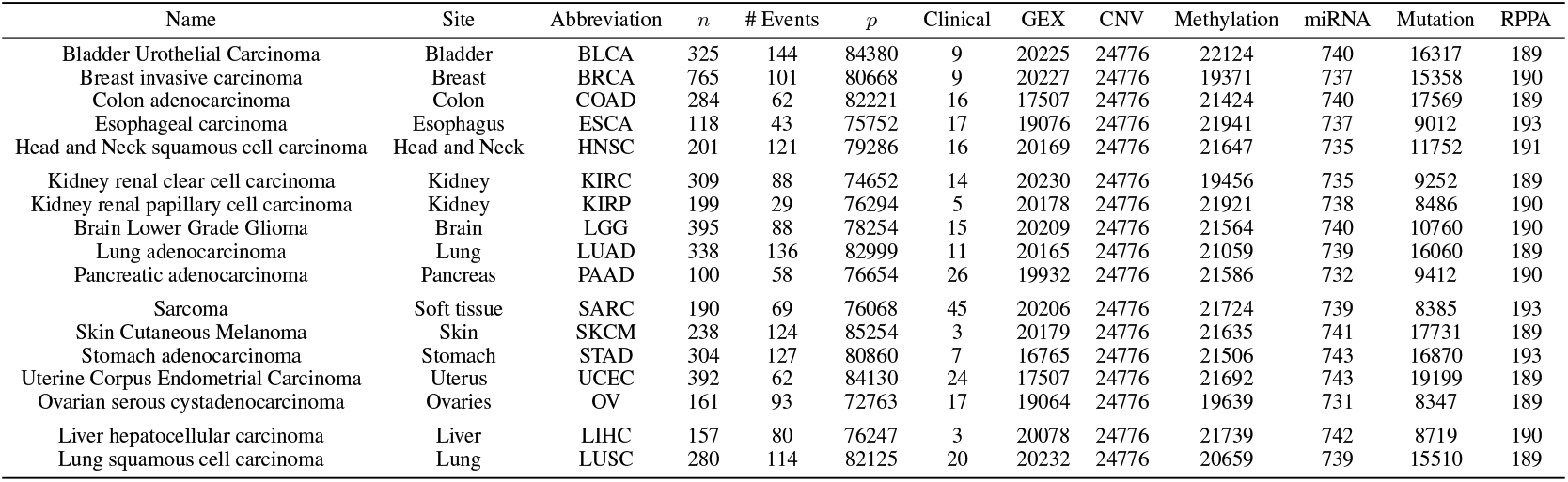
Summary information of all 17 considered TCGA datasets used in our study. High numbers of clinical features in some cancers are due to categorical features that were one-hot encoded. *n* denotes the number of samples per cancer. *p* denotes the number of total features per cancer. Modality names (*e*.*g*., Clinical) denote the number of features within each modality per cancer.

| Name | Site | Abbreviation | $n$ | # Events | $p$ | Clinical | GEX | CNV | Methylation | miRNA | Mutation | RPPA |
| --- | --- | --- | --- | --- | --- | --- | --- | --- | --- | --- | --- | --- |
| Bladder Urothelial Carcinoma | Bladder | BLCA | 325 | 144 | 84380 | 9 | 20225 | 24776 | 22124 | 740 | 16317 | 189 |
| Breast invasive carcinoma | Breast | BRCA | 765 | 101 | 80668 | 9 | 20227 | 24776 | 19371 | 737 | 15358 | 190 |
| Colon adenocarcinoma | Colon | COAD | 284 | 62 | 82221 | 16 | 17507 | 24776 | 21424 | 740 | 17569 | 189 |
| Esophageal carcinoma | Esophagus | ESCA | 118 | 43 | 75752 | 17 | 19076 | 24776 | 21941 | 737 | 9012 | 193 |
| Head and Neck squamous cell carcinoma | Head and Neck | HNSC | 201 | 121 | 79286 | 16 | 20169 | 24776 | 21647 | 735 | 11752 | 191 |
| Kidney renal clear cell carcinoma | Kidney | KIRC | 309 | 88 | 74652 | 14 | 20230 | 24776 | 19456 | 735 | 9252 | 189 |
| Kidney renal papillary cell carcinoma | Kidney | KIRP | 199 | 29 | 76294 | 5 | 20178 | 24776 | 21921 | 738 | 8486 | 190 |
| Brain Lower Grade Glioma | Brain | LGG | 395 | 88 | 78254 | 15 | 20209 | 24776 | 21564 | 740 | 10760 | 190 |
| Lung adenocarcinoma | Lung | LUAD | 338 | 136 | 82999 | 11 | 20165 | 24776 | 21059 | 739 | 16060 | 189 |
| Pancreatic adenocarcinoma | Pancreas | PAAD | 100 | 58 | 76654 | 26 | 19932 | 24776 | 21586 | 732 | 9412 | 190 |
| Sarcoma | Soft tissue | SARC | 190 | 69 | 76068 | 45 | 20206 | 24776 | 21724 | 739 | 8385 | 193 |
| Skin Cutaneous Melanoma | Skin | SKCM | 238 | 124 | 85254 | 3 | 20179 | 24776 | 21635 | 741 | 17731 | 189 |
| Stomach adenocarcinoma | Stomach | STAD | 304 | 127 | 80860 | 7 | 16765 | 24776 | 21506 | 743 | 16870 | 193 |
| Uterine Corpus Endometrial Carcinoma | Uterus | UCEC | 392 | 62 | 84130 | 24 | 17507 | 24776 | 21692 | 743 | 19199 | 189 |
| Ovarian serous cystadenocarcinoma | Ovaries | OV | 161 | 93 | 72763 | 17 | 19064 | 24776 | 19639 | 731 | 8347 | 189 |
| Liver hepatocellular carcinoma | Liver | LIHC | 157 | 80 | 76247 | 3 | 20078 | 24776 | 21739 | 742 | 8719 | 190 |
| Lung squamous cell carcinoma | Lung | LUSC | 280 | 114 | 82125 | 20 | 20232 | 24776 | 20659 | 739 | 15510 | 189 |

### 2.4 Reference models

For benchmarking purposes, we considered multiple reference models. In particular, we used the best performing model from the benchmark study of Herrmann et al. [2021], BlockForest. Further, we included RandomBlock favoring clinical features, a variant of BlockForest [Hornung and Wright, 2019a]. Favoring refers to RandomBlock mandatorily considering a particular group of variables, here clinical variables, in the split-point selection.

In addition, we included *prioritylasso* for which we favored (that is, did not penalize) clinical variables and used cross-validated offsets with five folds. The priority order was chosen to be (in descending order of priority) clinical, gene expression, mutation, miRNA, DNA methylation, copy number variation (CNV), and reversed-phase protein arrays (RPPA).

We also included a random survival forest method (RSF) and a Lasso regularized Cox PH model (Lasso) as two reference methods that did not use the group structure information present in the multi-omics variables. A Ridge regularized Cox PH model using only the clinical variables (clinical Cox PH) was also part of our benchmark. We opted for a Ridge regularized model since this allowed us to prevent convergence issues sometimes encountered even with dummy encoded categorical clinical variables.

RSF was implemented using *ranger* [Wright and Ziegler, 2017] *while Lasso and clinical Cox PH were implemented using glmnet* [Friedman et al., 2010, Simon et al., 2011]. *RandomBlock favoring clinical features and BlockForest were implemented using the BlockForest* package [Hornung and Wright, 2019b]. *prioritylasso* favoring clinical features was implemented using the *prioritylasso* package [Klau et al., 2020].

### 2.5 Datasets

We benchmarked all models on the TCGA datasets. Inspired by the approach of Herrmann et al. [2021], we selected cancers with at least 100 samples after preprocessing and an average event ratio greater or equal to five percent to ensure there were enough patients and events to calculate meaningful concordance values. Further, we used *GISTIC 2*.*0* CNV data [Mermel et al., 2011] and the number of non-silent mutation calls per gene per patient for mutation, the former of which was taken from Xena Browser [Goldman et al., 2020], with mutation coming from the PANCANATLAS [Chang et al., 2013]. Mutation calls were calculated from the PANCANATLAS MAF file using *Maftools* [Mayakonda et al., 2018]. In addition, we considered miRNA, mRNA, DNA methylation, RPPA, and clinical data, all of which were also taken from the PANCANATLAS. We log-transformed both mRNA and miRNA expression. Otherwise, no further preprocessing of the datasets was performed.

For comparability, we used the same clinical variables as Herrmann et al. [2021], with the caveat that we dropped clinical variables that were missing for more than five patients. Categorical clinical variables were one-hot encoded. We excluded omics variables if they were missing for more than one patient to preserve as many patients as possible. Table 1 shows an overview of all TCGA datasets which were used in our study.

### 2.6 Performance metric

Similar to other works [Cheerla and Gevaert, 2019, Tong et al., 2020, Hornung and Wright, 2019a], we used the c-index (Eq. 3) [Harrell et al., 1982] to measure the discriminative performance of our models, where 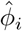 represents an estimated (relative) risk for patient *i*, and *T*_*i*_ = min(*Y*_*i*_, *S*_*i*_) represents the time until a patient was either censored or experienced the event. We have *δ* = 0 for censored patients and *δ* = 1 otherwise. Equivalently, the c-index is the ratio of concordant pairs and all comparable pairs.

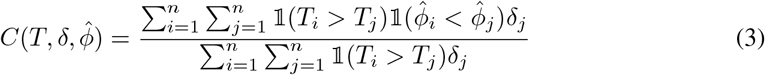

### 2.7 Validation

For all cancers in our dataset, we performed outer five-fold cross-validation, twice repeated, giving us a total of ten outer splits per cancer. Where applicable, we tuned the hyperparameters of each model using separate inner cross-validation with five folds or out-of-bag error (for random forest-based methods) to choose the best parameters to refit on each outer fold.

We used the *glmnet* internal *cv*.*glmnet* function to optimize the regularization parameter for clinical Cox PH and Lasso. BlockForest and RandomBlock favoring clinical features were tuned using the *blockForest::blockfor* function. For RSF, we tuned the *mtry* parameter using the *tuneRanger::tuneMtryFast* function.

For our neural networks, we fixed the *l*_2_ regularization hyperparameter *λ* at 0.001, since we were interested in comparing the learning dynamics of different integration methods with constant *l*_2_ regularization. In addition, research has shown that in the presence of batch normalization, *l*_2_ regularization amounts to changing the effective learning rate over time, which we already covered through learning rate decay, suggesting that *l*_2_ regularization should have a relatively negligible effect on our models [Hoffer et al., 2018, Van Laarhoven, 2017]. Early stopping and learning rate decay were used to implicitly control the number of epochs and the learning rate for all neural models, with an initial learning rate of 0.01.

13.968cm For statistical significance testing, we tested for an overall difference between models by adopting the same approach as Hornung and Wright [2019a]. We calculated the mean concordance per model per cancer (17 in total) and treated these mean values as independent between datasets. We then ran a one-sided paired t-test with a null hypothesis of non-inferiority of each reference method relative to the model of interest. The alternative hypothesis was that the model of interest performed significantly better than the respective benchmark method. In case any model failed to run on a test split, we imputed the missing test concordance(s) with the mean test concordance on the remaining splits.

## 3 Results

### 3.1 Hierarchical autoencoder-based integration outperforms other integration methods on multi-omics data

We found that HierarchicalSAE significantly outperformed all other considered integration techniques when integrating multi-omics data (Figure 2A). This pattern was consistent both when adding additional hidden layers to all of the considered integration methods (Figure S1) and when omitting the Cox supervision of the modality-specific autoencoders (Figure S2) (Table S1). Even though some of the other integration methods slightly improved their performance under the considered ablations, we consistently saw HierarchicalSAE decidedly outperform the other considered integration methods.

**Figure 2:**
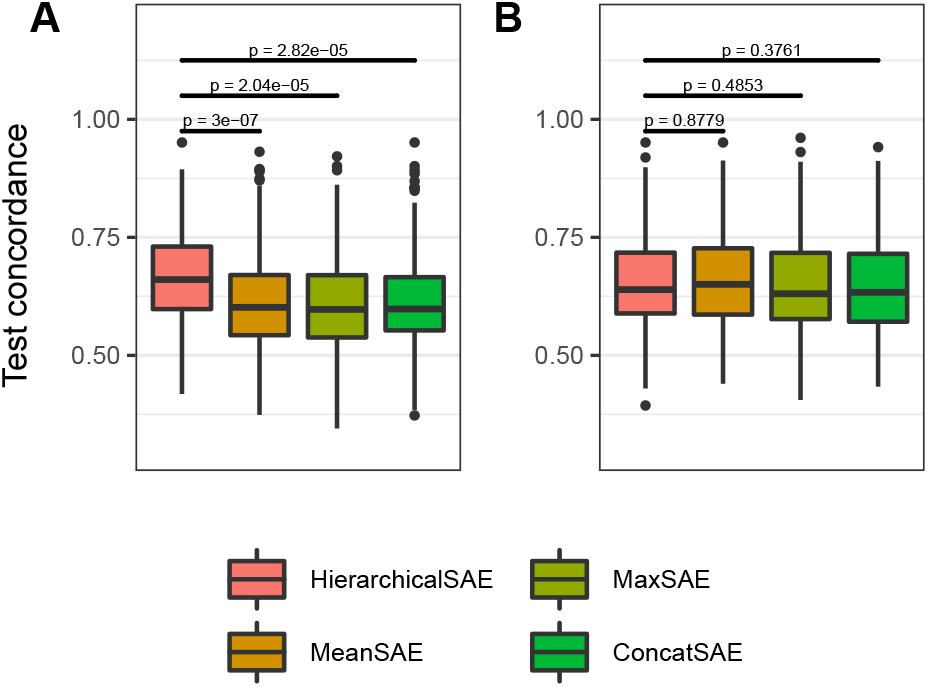
Performance of the four considered representational integration methods on TCGA. P-values testing for the superiority of HierarchicalSAE on bars. HierarchicalSAE outperforms other representational integration techniques with multi-omics data but not with clinical and gene expression. A: Overall test concordance across the 17 TCGA cancers when integrating all six omics blocks and clinical data. B. Overall test concordance across the 17 TCGA cancers when integrating gene expression and clinical data.

We considered two properties of our hierarchical autoencoder-based integration that may have contributed to its superior performance, relative to the other considered types of integration:

1. Forcing the final representation to be amenable to reconstructing (some of) the modality-specific representations may have made the final representation of HierarchicalSAE particularly well suited for survival prediction. The first property is thus focused only on the effect on the final representation as opposed to overall learning dynamics.
2. The additional regularization imposed by the reconstruction loss of the second-level autoencoder may have benefitted the performance of HierarchicalSAE. In particular, adding additional regularization through the reconstruction loss drove down the importance of the supervised loss(es), potentially changing overall learning dynamics.

To investigate these hypotheses for the outperformance of HierarchicalSAE, we first considered an ablation study in which we removed the decoder (and thus reconstruction loss) from HierarchicalSAE, which we refer to as HierarchicalSAE (no decoder) (Figure 1E). Removing the decoder affected both the learned representation directly (property one) and the overall level of regularization (property two). The performance of HierarchicalSAE (no decoder) model was decidedly worse than HierarchicalSAE and comparable to the other integration methods (Table 2).

**Table 2:** Performance of HierarchicalSAE (Figure 1D) under the considered ablations of no decoder (Figure 1E), increased *l*_2_ regularization hyperparameter *λ* and no early stopping. SD refers to the standard deviation across splits.

| Model | Ablation | Mean concordance (SD) |
| --- | --- | --- |
| HierarchicalSAE | None | 0.665 (0.104) |
| HierarchicalSAE | No early stopping (50 epochs) | 0.663 (0.106) |
| HierarchicalSAE | No decoder + higher $\ell_2$ regularization ($\lambda = 0.1$) | 0.628 (0.107) |
| HierarchicalSAE | No decoder + higher $\ell_2$ regularization ($\lambda = 0.01$) | 0.625 (0.101) |
| HierarchicalSAE | No decoder + no early stopping | 0.609 (0.095) |
| HierarchicalSAE | No decoder | 0.608 (0.099) |

Next, we increased the regularization of HierarchicalSAE (no decoder) by increasing its *l*_2_ regularization hyperparameter *λ*, to inform whether removing the decoder decreased performance due to a lower overall of regularization, relative to HierarchicalSAE. In addition to the default value of *λ* = 0.001, we also considered *λ* = 0.01 and *λ* = 0.1. While increasing the hyperparameter *λ* helped HierarchicalSAE (no decoder) slightly, its performance was still not comparable to that of HierarchicalSAE (Table 2), suggesting that the increased overall regularization (property two) of HierarchicalSAE was not primarily responsible for its outperformance.

As a final ablation, we ran both HierarchicalSAE and HierarchicalSAE (no decoder) with early stopping turned off, to investigate whether the addition or omission of the reconstruction loss was causing either of the models to train especially long or short (property two). Since we noticed that our models would often stop before their maximum of 100 epochs, we ran both models for a fixed 50 epochs in this ablation. Similar to the other ablations, turning off early stopping did not noticeably close the gap between HierarchicalSAE and HierarchicalSAE (no decoder) (Table 2). Since neither increasing *l*_2_ regularization nor turning off early stopping closed the gap between HierarchicalSAE and HierarchicalSAE (no decoder), we concluded that the reconstruction loss enabled HierarchicalSAE to learn an integrated representation that was particularly effective for multi-omics integration.

We also compared the performance of all integration techniques when integrating only clinical data and gene expression, an arguably much simpler task, on which many multi-omics models perform as good or better than when integrating all available modalities [Hornung and Wright, 2019a, Vale-Silva and Rohr, 2021]. When trained on only clinical and gene expression data, the gap between integration methods disappeared (Figure 2B). This suggested that simpler integration techniques (such as mean-pooling, max-pooling, and concatenation) might be affected more by the noise imposed by adding further (potentially less informative) modalities. HierarchicalSAE, on the other hand, was better able to handle additional modalities, performing better when integrating all available modalities compared to integrating clinical and gene expression only, a pattern which was reversed for all other considered representational integration methods (Figure 2).

### 3.2 Hierarchical autoencoder-based integration performs on-par with the existing state of the art statistical multi-omics methods

Overall, HierarchicalSAE significantly outperformed the clinical Cox PH model (*p* = 0.0146), slightly outperformed BlockForest (although the difference was not statistically significant), and was comparable with RandomBlock favoring clinical features (Figure 3 and Table 3). That said, RandomBlock favoring clinical features had the advantage of favoring clinical variables, that is, the model was *a priori* provided with the knowledge that clinical data contained strong survival-relevant information. HierarchicalSAE was not provided with information about the importance of clinical data, yet performed competitively with RandomBlock favoring clinical features overall and for cancers on which clinical data was informative. MaxSAE, ConcatSAE, and MeanSAE all performed comparable to the Lasso, suggesting that they were effectively not able to perform multi-omics integration since the Lasso treats all modalities uniformly (Table 3).

**Table 3:** Mean test concordance pooled across the 17 considered TCGA datasets. Clinical Cox PH included as a baseline model that uses only clinical data in both settings. P-values correspond to testing non-inferiority to HierarchicalSAE using a one-sided paired t-test. Best value is shown in bold. SD refers to the standard deviation across splits.

| Model | Multi-omics data |  | Clinical and gene expression |  |
| --- | --- | --- | --- | --- |
|  | Mean concordance (SD) | P-value | Mean concordance (SD) | P-value |
| HierarchicalSAE | **0.665 (0.104)** | - | 0.652 (0.109) | - |
| RandomBlock favoring | 0.663 (0.113) | 0.42179 | 0.652 (0.113) | 0.49704 |
| BlockForest | 0.649 (0.118) | 0.07653 | **0.663 (0.113)** | 0.92535 |
| prioritylasso favoring | 0.641 (0.119) | 0.01344 | 0.641 (0.117) | 0.08647 |
| MeanSAE | 0.617 (0.109) | < 1e-06 | 0.662 (0.109) | 0.8779 |
| Lasso | 0.614 (0.12) | 0.00044 | 0.623 (0.119) | 0.00775 |
| ConcatSAE | 0.613 (0.104) | 3e-05 | 0.65 (0.11) | 0.37614 |
| MaxSAE | 0.61 (0.112) | 2e-05 | 0.652 (0.111) | 0.48532 |
| RSF | 0.595 (0.125) | 9e-05 | 0.61 (0.117) | 0.00512 |
| Clinical Cox PH (baseline) | 0.622 (0.112) | 0.01457 | 0.622 (0.112) | 0.02378 |

**Figure 3:**
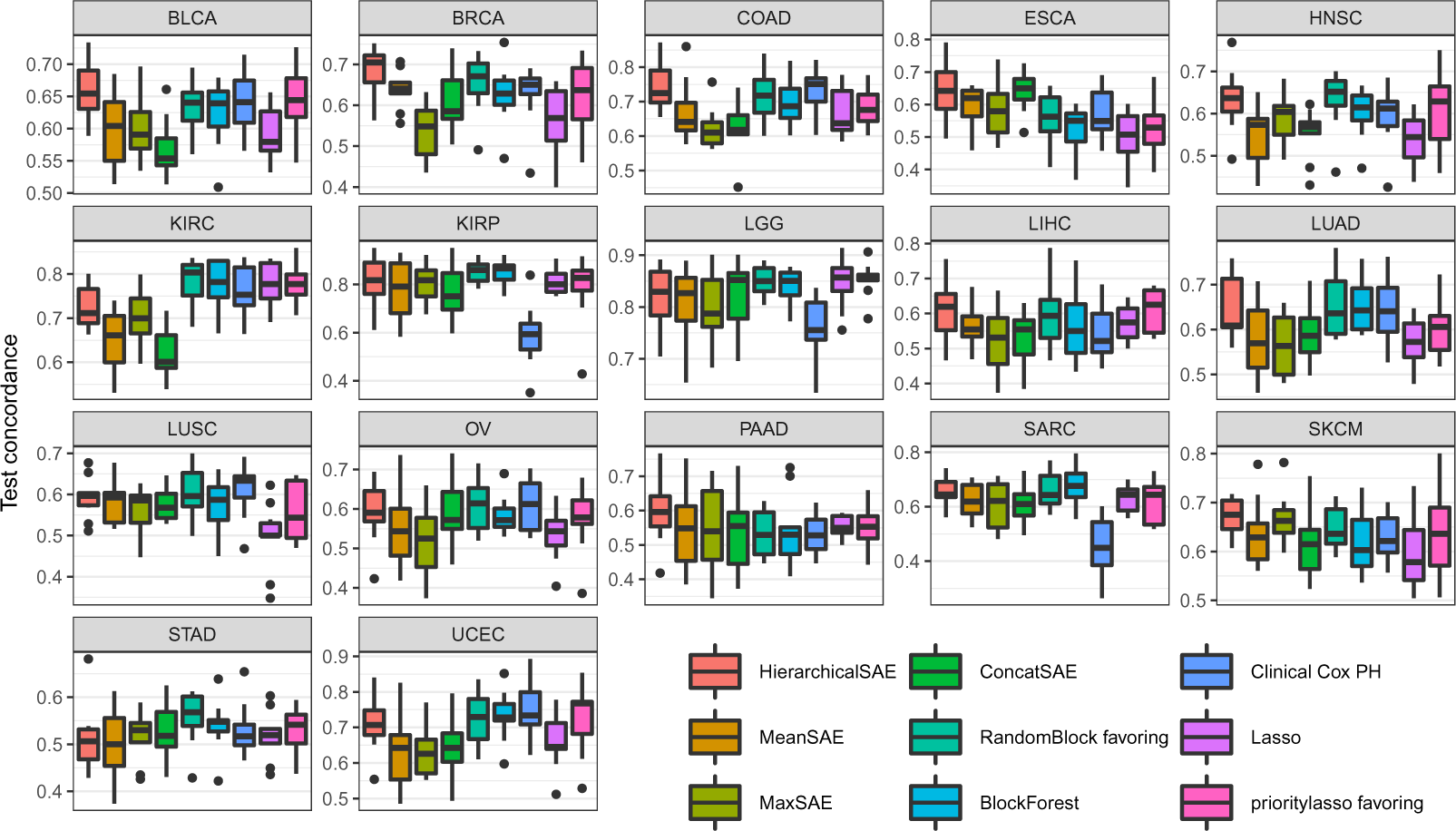
Per cancer test concordance of all models on each of the 17 TCGA cancers when integrating six omics blocks and clinical data. We observed strong hetereoneity in terms of which model performed best across different cancers.

We observed strong heterogeneity in terms of which model performed best across datasets. Both RandomBlock favoring clinical features and HierarchicalSAE achieved the highest mean concordance out of all models on six out of the 17 considered datasets, with *prioritylasso* favoring clinical features, the clinical Cox PH model and BlockForest also performing best in terms of mean concordance on at least one cancer (Figure 3 and Table S2). In addition, we found that on most cancers, the clinical Cox PH model was competitive with the multi-omics methods, with a few exceptions such as SARC, KIRP, and LGG, which led to the overall underperformance of the Cox PH model.

Lastly, our results once again confirmed the pattern often seen in multi-omics studies, namely that many methods perform better with only a subset of modalities. Specifically, only RandomBlock favoring clinical features and HierarchicalSAE were able to improve their performance when integrating additional modalities beyond clinical data and gene expression, while *prioritylasso* favoring clinical features maintained its overall performance. All other models produced worse performance when attempting to integrate all seven modalities compared to only using clinical and gene expression data, some strongly so (Table 3). This observation motivates the question of why HierarchicalSAE and RandomBlock favoring clinical features were able to improve their performance while all other methods struggled to do so. We leave the exploration of this question for RandomBlock favoring clinical features to future work and focus on HierarchicalSAE.

### 3.3 Multi-omics integration as group-wise feature selection

To better understand the representations learned by HierarchicalSAE, we first compared the centered kernel alignment (CKA) similarities [Kornblith et al., 2019] of all modality-specific representations, as well as that of the final layer before the predicted risk for HierarchicalSAE and HierarchicalSAE (no decoder) for selected cancers. Kornblith et al. [2019] proposed to use CKA as a metric to measure similarity between neural network representations that is robust to initialization, invariant to invertible linear transformations, and can “measure meaningful similarities between representations of higher dimension than the number of data points” [Kornblith et al., 2019].

We chose to focus on bladder cancer (BLCA) and sarcoma (SARC) as HierarchicalSAE outperformed the version without a decoder strongly in both cases (Table S2). In addition, the clinical Cox PH model performed quite competitively on BLCA, but underperformed on SARC (Figure 3), making these cancers useful also to further study the difference between our multi-omics method and a clinical-only model.

We found that the modality-specific representation of HierarchicalSAE and HierarchicalSAE (no decoder) were all reasonably similar to each other (*e*.*g*., clinical was similar to clinical on TCGA-BLCA), with CKA similarities greater than 0.5. At the final representation layer, however, we observed lower CKA similarity between the two architectures, presumably caused by the omission of the reconstruction loss for the version of HierarchicalSAE without a decoder (Figure 4). In addition, we observed that while the final representation of HierarchicalSAE was not similar to the final representation of HierarchicalSAE (no decoder), it was similar in both cancers to at least one of the modality-specific representations of HierarchicalSAE (no decoder), suggesting that the reconstruction loss might be causing HierarchicalSAE to focus on selected modalities. For BLCA, where the clinical Cox PH model performed well, the final representation of HierarchicalSAE was strongly similar to the modality-specific representation of the clinical block of HierarchicalSAE (no decoder). For SARC, on the other hand, where the clinical Cox PH model underperformed, the final representation had its strongest CKA similarities primarily with miRNA and other omics modalities.

**Figure 4:**
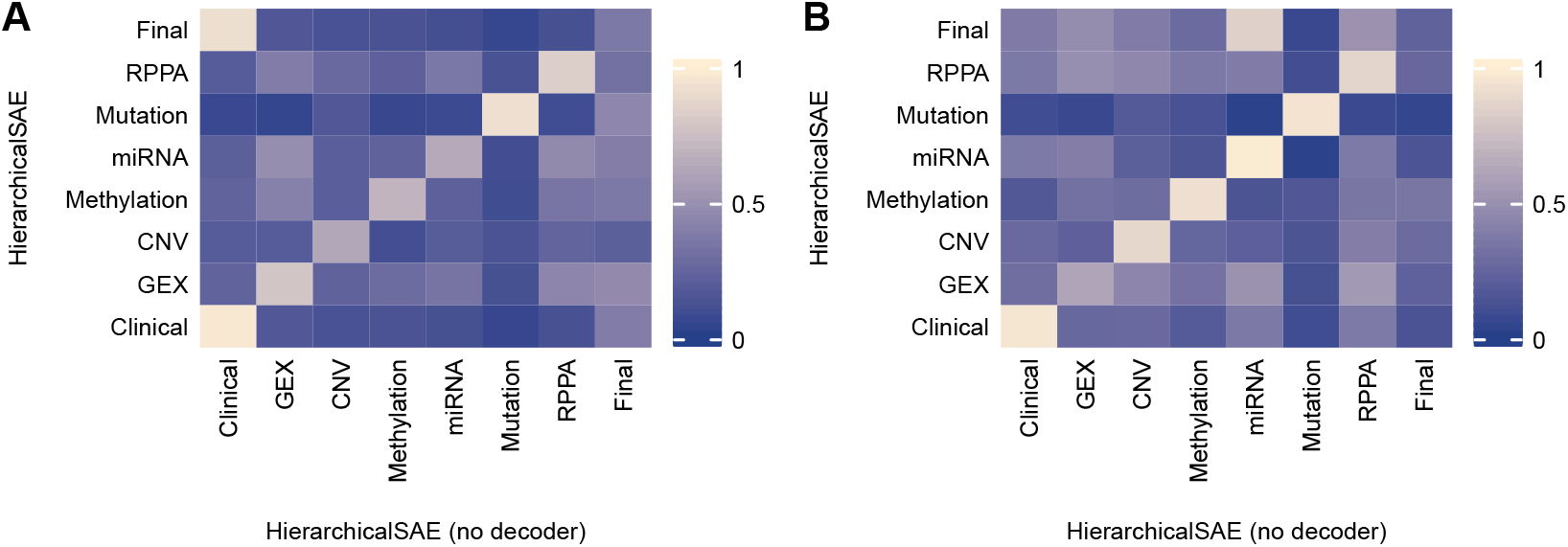
CKA representational similarities between the representations of each modality-specific autoencoder and the final representation before the risk prediction of HierarchicalSAE and HierachicalSAE (no decoder). HierarchicalSAE learned final representations that were strongly similar to one or more of the modality-specific representations of HierarchicalSAE (no decoder), see the first row and right-most column. A. CKA representational similarities on the first test split of TCGA-BLCA. B. CKA representational similarities on the first test split of TCGA-SARC.

Motivated by the observation that HierarchicalSAE tended to focus on modalities potentially important for each cancer, we ran an auxiliary experiment in which we pruned modalities from HierarchicalSAE and HierarchicalSAE (no decoder) after concatenation in a specific order and observed any performance changes without retraining (Figure S4. We considered two pruning orders, first, *l*_1_ magnitude pruning, for which we first pruned the modality with the lowest *l*_1_ norm of its edges after concatenation of all modality-specific representations. Secondly, we considered CKA pruning, in which we first pruned the modality whose modality-specific representation exhibited the lowest CKA similarity with the final representation of the considered model. We found that under both considered modality orderings, the performance of HierarchicalSAE relative to no pruning degraded more slowly than that of HierarchicalSAE (no decoder) (Figure 5). This strengthened our hypothesis that the reconstruction loss of the hierarchical autoencoder-based integration was forcing HierarchicalSAE to focus on important modalities. Moreover, when considering absolute performance changes as modality pruning was performed, we found that HierarchicalSAE decreased its performance linearly with the pruning of each modality (Figure S3), suggesting that modalities were contributing additively, that is, that there were no major interactions between modalities. In particular, under magnitude pruning, the median performance of HierarchicalSAE with two modalities remaining (*i*.*e*., having pruned five modalities) was equal to the median performance when training HierarchicalSAE on clinical and gene expression from scratch (Figure S3A). For HierarchicalSAE (no decoder), on the other hand, its performance was much worse with all considered modalities than with only clinical and gene expression, and modality pruning only widened this gap (Figure S3A-B).

**Figure 5:**
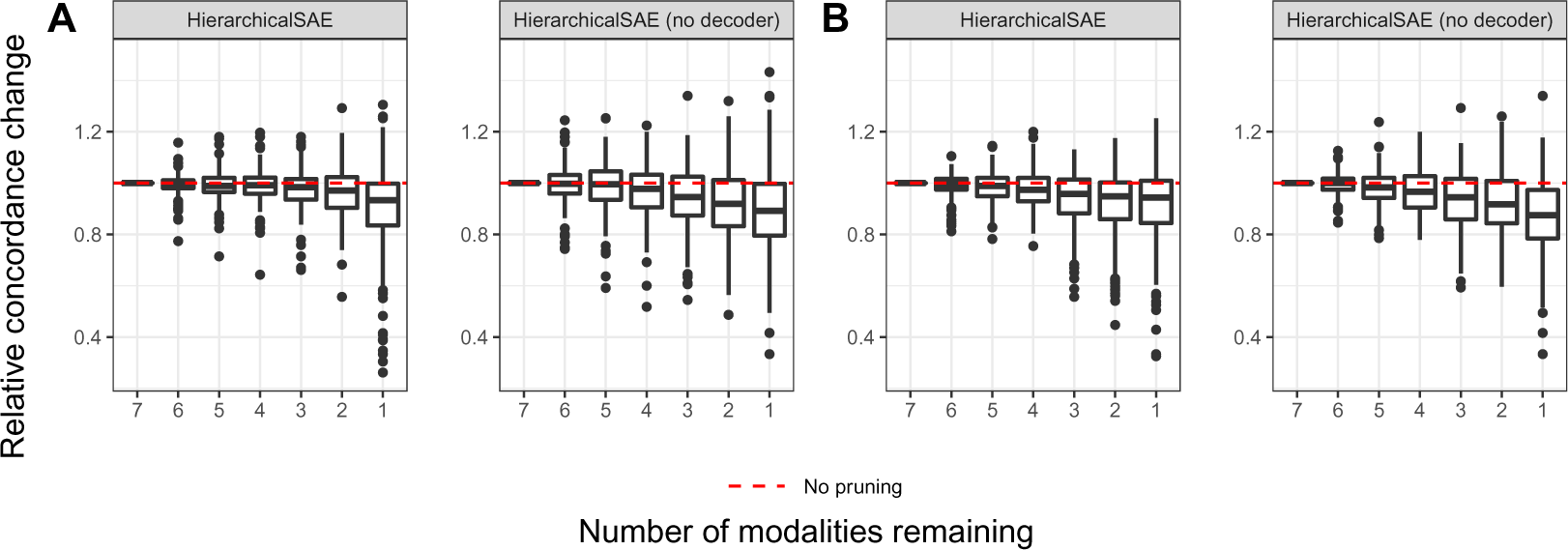
Performance changes of HierarchicalSAE and HierarchicalSAE (no decoder) as modalities were pruned in order without retraining, relative to no pruning (Figure S4). A. Performance under CKA modality pruning in decreasing order of CKA similarity with the final representation. B. Performance under *l*_1_ magnitude pruning based on the total *l*_1_ norm of each modality after concatenation.

## 4 Discussion

We have shown that hierarchical autoencoder-based integration was more effective than other techniques for integrating modality-specific representations in a multi-omics context. Our goal within this work was to critically evaluate different methods for representational integration, with a focus on why methods tended to work well. Our results indicated that well-performing multi-omics integration (at least within a survival context) implies not necessarily the integration of all modalities but rather also potential group-level feature selection. Our modality selection hypothesis is supported by the fact nearly all models performed better with only a subset of available modalities (Table 3).

In addition, our work showed that even methods that do work with multi-omics data, such as HierarchcialSAE, likely do so partially because they are implicitly focusing on the most important modalities. That said, both RandomBlock favoring clinical features and HierarchicalSAE performed better with all multi-omics data than with only clinical and gene expression, suggesting that perhaps these models were able to learn additional information from the added modalities, even if they only slightly outperformed the best model trained on clinical and gene expression only (Table 3).

More broadly, we established that hierarchical autoencoder-based representational integration could be leveraged to achieve competitive performance on the TCGA dataset as measured by the c-index. We believe our benchmark is especially interesting since there has not been a conclusive comparison of neural models for multi-omics integration to state-of-the-art statistical methods such as BlockForest. Most previous work has tended to either focus on statistical methods only [Herrmann et al., 2021] or did not include dedicated multi-omics methods such as BlockForest as statistical reference methods. Our results thus validated hierarchical autoencoder-based representational integration as a solid alternative to BlockForest and *prioritylasso* favoring clinical features. This holds especially given the fact that RandomBlock and *prioritylasso* favoring clinical features had access to the *a priori* information that clinical data was important on TCGA and we gave *prioritylasso* additional information in the form of our selected priority order.

Our results were mostly coherent with the benchmarking study of Herrmann et al. [2021]. One difference worth mentioning was the fact that in our study, the clinical-only Cox PH model underperformed, compared to the work of Herrmann et al. [2021], relatively speaking. We believe this worsened performance was caused primarily by the fact that we did not include all clinical features that Herrmann et al. [2021] did, as we wanted to keep as many samples as possible. That said, our study was *not* meant not to rigorously establish the additional predictive value of multi-omics models beyond clinical. Other studies such as Vale-Silva and Rohr [2021] have also found additional predictive value of omics beyond clinical, but the (additional) predictive value of multi-omics beyond clinical is likely strongly dependant on both the cancer type in question and the clinical variables used. Sensitivity analyses varying the threshold at which clinical features with high missingness would be excluded would be useful to further investigate the trade-off between keeping more patients and using a higher number of clinical variables. That said, in our study, still most models underperformed the clinical Cox PH model overall, with only a few multi-omics integration methods consistently outperforming it. We also used a different type of concordance for measuring discriminative performance, which may have caused further difference relative to Herrmann et al. [2021].

There are various promising avenues of future work connected with our approach in this study that we did not pursue further. First, it could be interesting to evaluate the performance of different methods (both multi-omics and naive) while explicitly restricting the number of usable modalities. Future research might focus on attempting to find the most predictive modalities (even if these are known to be clinical and gene expression *a priori*, good models should be able to find them) as opposed to explicitly trying to integrate all available ones. An alternative approach might be vertical multi-omics integration, in which modalities are first integrated at the gene level after which they are again integrated to find a representation amenable to a final survival or risk prediction. While there has been some work on this topic, there is a need to further explore this direction, including exhaustively benchmarking vertical multi-omics integration against standard multi-omics integration techniques such as BlockForest.

Our work included some limitations. One limitation is that we only benchmarked our models on TCGA; moreover, we only considered a total of 17 out of the 33 available cancers due to concerns of low sample size or missing omics modalities in the remaining datasets. Even though this is a common strategy among researchers [Herrmann et al., 2021, Hornung and Wright, 2019a, Cheerla and Gevaert, 2019], future work should run a comparable benchmark, including statistical methods and neural networks on another large-scale multi-omics survival dataset. In addition, we did not include pan-cancer-based methods in our benchmarks, as we were focused on establishing well-working integration methods in the simpler case of single-cancer training.

## 5 Conclusion

Our work demonstrated that hierarchical autoencoder-based integration was effective for multi-omics integration in cancer survival models as benchmarked on the TCGA datasets. The models we trained using hierarchical autoencoder-based integration performed as well or better than the best statistical methods reported in a recent large-scale benchmarking study [Herrmann et al., 2021], BlockForest, and one of its variants, RandomBlock favoring clinical features.

Moreover, we showed that the specific integration method chosen for representational integration of modality-specific neural networks mattered very little when the right modalities were chosen *a priori*. When all available modalities were to be integrated, however, hierarchical autoencoder-based integration strongly outperformed other integration techniques such as max-pooling. Based on this finding, we also posited that multi-omics integration was partially reducible to a group-wise feature selection problem. In particular, we showed that our hierarchical autoencoder-based integration performed well partially because it selected out the most informative modalities on each cancer type (as opposed to integrating every available modality).

## Supporting information

Full Supplement

## Acknowledgements

The results shown here are in whole or part based upon data generated by the TCGA Research Network: https://www.cancer.gov/tcga. Benchmarks were run on the Euler cluster managed by the HPC team at ETH Zurich.

