## Supplementary material for "Hierarchical autoencoder-based integration improves performance in multi-omics cancer survival models through soft modality selection": Full Supplement

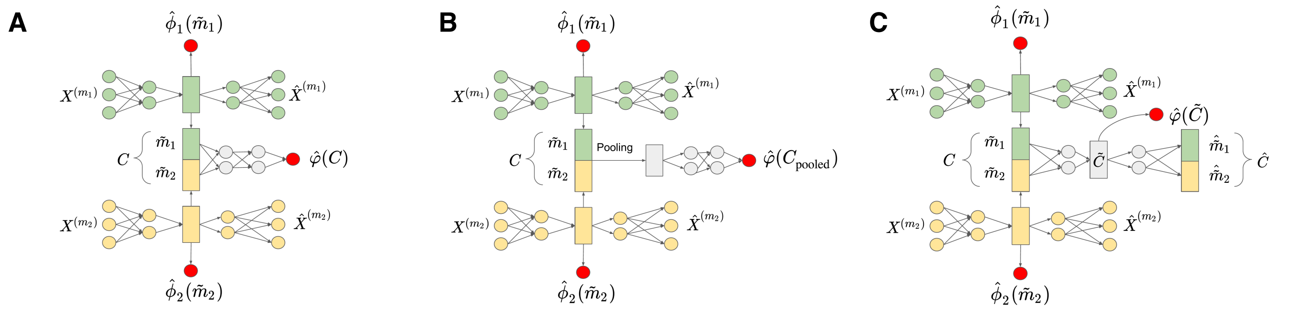


**Fig. S1.** Architecture diagrams of all considered representational integration methods with additional hidden layers. The number of hidden layers for ConcatSAE, MeanSAE, and MaxSAE was chosen to mirror the architecture of HierarchicalSAE. Non-linear activations were added only after the first hidden layer. Two input modalities shown for simplicity. Let $X^{(m_i)}$ denote the submatrix of $X$ containing only columns belonging to the i-th input variable group, let $\hat{X}^{(m_i)}$ denote a reconstruction of $X^{(m_i)}$. Let $\tilde{m}_i$ be the modality-specific representation learned based on input modality $i$. Let $\hat{\phi}_1, \ldots, \hat{\phi}_{|M|}$ be predicted relative risks based on the modality-specific representation learned for each input modality, while $\hat{\varphi}$ represents the predicted relative risk based on the overall joint representation. A. Architecture diagram of ConcatSAE with additional hidden layers after the integration. B. Architecture diagram of MeanSAE and MaxSAE with additional hidden layers after the integration. Pooling may refer to either max-pooling or mean-pooling. C. Architecture diagram of HierachicalSAE with an additional hidden layer as part of the integration.

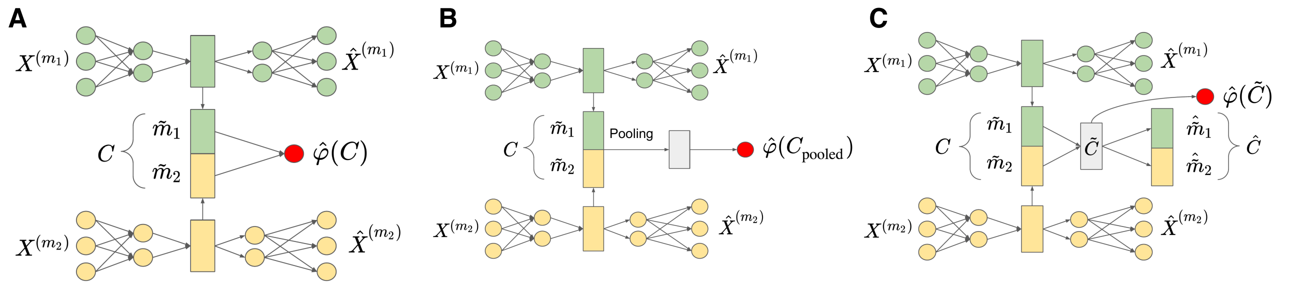


**Fig. S2.** Architecture diagrams of all considered representational integration methods without supervision of the modality specific autoencoders. Two input modalities shown for simplicity. Let $X^{(m_i)}$ denote the submatrix of $X$ containing only columns belonging to the i-th input variable group, let $\hat{X}^{(m_i)}$ denote a reconstruction of $X^{(m_i)}$. Let $\tilde{m}_i$ be the modality-specific representation learned based on input modality $i$. Let $\hat{\varphi}$ represent the predicted relative risk based on the overall joint representation. A. Architecture diagram of ConcatSAE without supervision. B. Architecture diagram of MeanSAE and MaxSAE without supervision. Pooling may refer to either max-pooling or mean-pooling. C. Architecture diagram of HierachicalSAE without supervision.

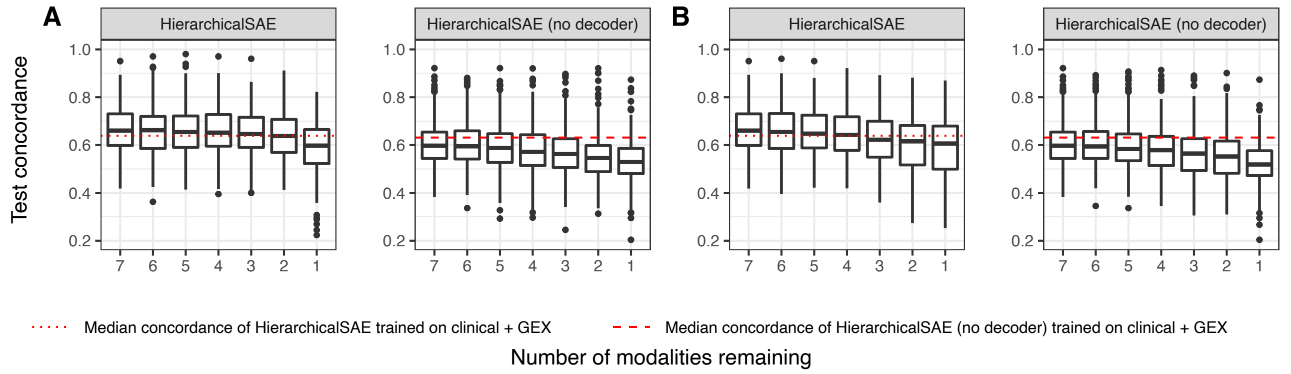


**Fig. S3.** Absolute performance changes of HierarchicalSAE and HierarchicalSAE (without deocder) as full modalities were pruned in order without retraining (Fig. S4). A. Performance under CKA modality pruning in decreasing order of CKA similarity with the final representation. B. Performance under $\ell_1$ magnitude pruning based on total $\ell_1$ norm of each modality after the concatenation..

Table S1. Mean concordance pooled across all 17 TCGA datasets of all considered representational integration methods without any ablations (Fig. 1B-D), with additional hidden layers (Fig. S1A-C) and without supervision of the modality-specific autoencoders (Fig. S2A-C). SD refers to the standard deviation across splits.

| Model | Ablation | Mean concordance (SD) |
|---|---|---|
| HierarchicalSAE | None | 0.665 (0.104) |
| HierarchicalSAE | Additional hidden layer | 0.658 (0.109) |
| HierarchicalSAE | No supervision of modality-specific autoencoders | 0.662 (0.108) |
| MeanSAE | None | 0.617 (0.109) |
| MeanSAE | Additional hidden layer | 0.619 (0.107) |
| MeanSAE | No supervision of modality-specific autoencoders | 0.631 (0.11) |
| MaxSAE | None | 0.61 (0.112) |
| MaxSAE | Additional hidden layer | 0.601 (0.114) |
| MaxSAE | No supervision of modality-specific autoencoders | 0.608 (0.113) |
| ConcatSAE | None | 0.613 (0.104) |
| ConcatSAE | Additional hidden layer | 0.623 (0.107) |
| ConcatSAE | No supervision of modality-specific autoencoders | 0.629 (0.1) |

Table S2. Mean concordance across test splits for each of the 17 considered TCGA cancer datasets for all considered models rounded to three digits for display purposes. Best value by cancer in bold. Standard deviation across splits in parantheses.

| Model | BLCA | BRCA | COAD | ESCA | HNSC | KIRC | KIRP | LGG | LIHC | LUAD | LUSC | OV | PAAD | SARC | SKCM | STAD | UCEC |
|---|---|---|---|---|---|---|---|---|---|---|---|---|---|---|---|---|---|
| BlockForest | 0.624 (0.052) | 0.626 (0.073) | 0.696 (0.07) | 0.525 (0.075) | 0.602 (0.059) | 0.779 (0.055) | 0.852 (0.05) | 0.84 (0.035) | 0.566 (0.103) | 0.651 (0.058) | 0.573 (0.07) | 0.584 (0.046) | 0.539 (0.104) | **0.678 (0.071)** | 0.62 (0.067) | 0.541 (0.054) | 0.731 (0.07) |
| clinical Cox PH | 0.642 (0.045) | 0.629 (0.074) | 0.731 (0.063) | 0.575 (0.078) | 0.594 (0.07) | 0.76 (0.061) | 0.59 (0.129) | 0.759 (0.063) | 0.542 (0.079) | 0.641 (0.075) | **0.614 (0.068)** | 0.61 (0.068) | 0.533 (0.058) | 0.453 (0.112) | 0.629 (0.051) | 0.528 (0.057) | **0.752 (0.083)** |
| ConcatSAE | 0.568 (0.046) | 0.608 (0.07) | 0.628 (0.08) | 0.64 (0.06) | 0.554 (0.059) | 0.62 (0.053) | 0.768 (0.109) | 0.819 (0.075) | 0.526 (0.082) | 0.589 (0.067) | 0.576 (0.041) | 0.596 (0.089) | 0.531 (0.108) | 0.609 (0.068) | 0.613 (0.07) | 0.528 (0.056) | 0.646 (0.085) |
| HierarchicalSAE | **0.661 (0.046)** | **0.681 (0.067)** | **0.746 (0.074)** | **0.646 (0.095)** | 0.634 (0.074) | 0.725 (0.049) | 0.814 (0.099) | 0.819 (0.063) | 0.611 (0.087) | 0.646 (0.074) | 0.593 (0.05) | 0.593 (0.08) | **0.596 (0.093)** | 0.653 (0.06) | **0.672 (0.036)** | 0.512 (0.07) | 0.71 (0.077) |
| HierarchicalSAE (no decoder) | 0.576 (0.049) | 0.614 (0.072) | 0.626 (0.069) | 0.604 (0.067) | 0.543 (0.058) | 0.606 (0.042) | 0.753 (0.097) | 0.807 (0.064) | 0.538 (0.075) | 0.592 (0.056) | 0.585 (0.052) | 0.568 (0.073) | 0.544 (0.106) | 0.608 (0.079) | 0.607 (0.077) | 0.526 (0.043) | 0.644 (0.105) |
| Lasso | 0.593 (0.044) | 0.556 (0.094) | 0.667 (0.069) | 0.505 (0.08) | 0.54 (0.064) | 0.778 (0.05) | 0.812 (0.057) | 0.847 (0.048) | 0.575 (0.051) | 0.574 (0.053) | 0.497 (0.082) | 0.535 (0.064) | 0.553 (0.033) | 0.631 (0.05) | 0.596 (0.072) | 0.518 (0.051) | 0.658 (0.077) |
| MaxSAE | 0.599 (0.046) | 0.537 (0.067) | 0.621 (0.059) | 0.58 (0.083) | 0.592 (0.057) | 0.702 (0.068) | 0.807 (0.079) | 0.796 (0.068) | 0.525 (0.096) | 0.564 (0.071) | 0.565 (0.054) | 0.518 (0.086) | 0.545 (0.125) | 0.605 (0.091) | 0.664 (0.054) | 0.516 (0.051) | 0.628 (0.071) |
| MeanSAE | 0.6 (0.059) | 0.639 (0.046) | 0.67 (0.088) | 0.598 (0.063) | 0.546 (0.07) | 0.645 (0.075) | 0.779 (0.124) | 0.806 (0.075) | 0.561 (0.056) | 0.578 (0.086) | 0.584 (0.058) | 0.55 (0.096) | 0.545 (0.115) | 0.624 (0.064) | 0.638 (0.068) | 0.501 (0.071) | 0.635 (0.105) |
| *prioritylasso* favoring clinical features | 0.643 (0.057) | 0.617 (0.1) | 0.679 (0.056) | 0.531 (0.092) | 0.61 (0.093) | 0.778 (0.042) | 0.788 (0.141) | **0.854 (0.033)** | **0.611 (0.064)** | 0.602 (0.061) | 0.557 (0.073) | 0.572 (0.081) | 0.55 (0.059) | 0.617 (0.079) | 0.637 (0.085) | 0.53 (0.052) | 0.725 (0.096) |
| RandomBlock favoring clinical features | 0.633 (0.041) | 0.655 (0.07) | 0.717 (0.071) | 0.552 (0.087) | **0.634 (0.07)** | **0.784 (0.05)** | **0.853 (0.046)** | 0.851 (0.03) | 0.603 (0.108) | **0.655 (0.075)** | 0.602 (0.064) | **0.613 (0.068)** | 0.535 (0.069) | 0.66 (0.071) | 0.649 (0.044) | **0.557 (0.057)** | 0.724 (0.079) |

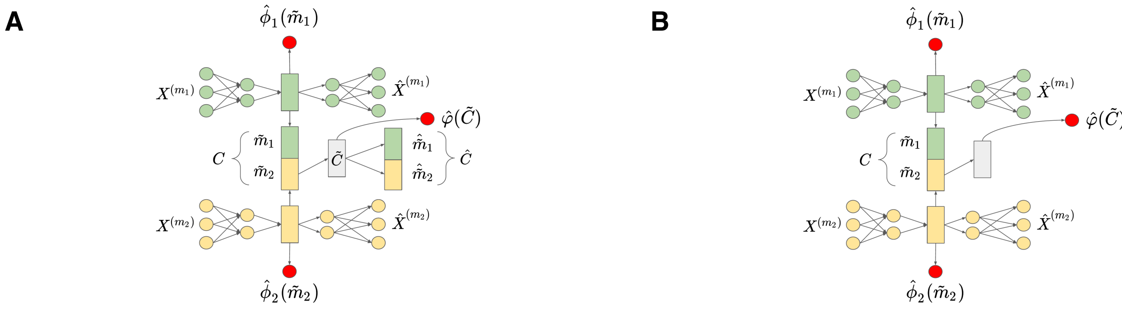


**Fig. S4.** Architecture diagrams of HierarchicalSAE and HierarchicalSAE (no decoder) with modality pruning. Two input modalities shown for simplicity. Let $X^{(m_i)}$ denote the submatrix of $X$ containing only columns belonging to the i-th input variable group, let $\hat{X}^{(m_i)}$ denote a reconstruction of $X^{(m_i)}$. Let $\tilde{m}_i$ be the modality-specific representation learned based on input modality $i$. Let $\hat{\phi}_1, \ldots, \hat{\phi}_{|M|}$ be predicted relative risks based on the modality-specific representation learned for each input modality, while $\hat{\varphi}$ represents the predicted relative risk based on the overall joint representation. A. Architecture diagram of HierarchicalSAE with modality $m_1$ pruned. B. Architecture diagram of HierarchicalSAE (no decoder) with modality $m_1$ pruned.
